# Evidence for an apathy phenotype in aged mice

**DOI:** 10.1101/2020.05.28.121004

**Authors:** Megan G Jackson, Stafford L Lightman, Gary Gilmour, Hugh Marston, Emma S J Robinson

## Abstract

Apathy is widely reported in patients with neurological disorders or post viral infection but is also seen in otherwise-healthy aged individuals. This study investigated whether aged mice express behavioural and physiological changes indicative of an apathy phenotype. Using measures of motivation to work for reward, we found deficits in the progressive ratio task related to rate of responding. In an effort for reward task, aged mice were less willing to exert effort for high value reward. Aged mice exhibited reduced reward sensitivity and expressed lower measures of anxiety in the novelty supressed feeding test. In a test of cognition (novel object recognition) aged mice showed no impairments but activity was lower in a measure of exploration in a novel environment. Aged mice also showed an attenuated response to restraint stress with lower corticosterone and reduced paraventricular nucleus c-fos activation. Together, these data suggest aged mice show reduced goal-directed behaviour and reduced reward sensitivity and stress reactivity, reflective of emotional blunting and may be a suitable model for pre-clinical apathy research.

## 1. Introduction

Aging is associated with widespread physiological changes, including disruption to normal behaviour. A prevalent behavioural change is the onset of the clinical symptom, apathy. Apathy is defined as a quantitative reduction in self-generated or voluntary behaviours (Levy and Czernecki, 2006). While apathy is a common feature of neurodegenerative diseases, including Alzheimer’s disease and Parkinson’s disease, and in some people following viral infections (Kamat et al., 2012), it is also seen in otherwise healthy aging. Apathy has a profound effect on the daily functioning of the individual. It is associated with cognitive decline, nutritional deficit and an overall poorer quality of life (Gerritsen et al., 2005; Ishii et al., 2009). Beyond the impact to the individual, it also significantly increases stress of the family and caregivers.

While generally considered a motivational disorder, the work of Marin, Levy and Dubois has conceptualised apathy into three domains. An emotional/affective component, where the individual can no longer link emotional signals with behaviour, an auto-activation/behavioural component where the individual can no longer self-initiate actions, and a cognitive component, where the individual lacks the ability to form an idea, and loses curiosity and routine (Levy and Dubois, 2006). Efforts have been made to map these components onto distinct brain circuits, particularly those of the frontal cortex-basal ganglia (Levy and Dubois, 2006) although there have been few studies to understand the cause of apathy in the context of normal aging. The development of a suitable method to model apathy in non-human species could facilitate insight into its underlying neurobiology and potentially elucidate therapeutic targets. As such, it is a crucial first step to establish whether aged rodents display behaviours relevant to the apathy domains seen in the human literature.

Clinical assessment of apathy is traditionally conducted using self-report or questionnaire-based methods (e.g. Apathy Evaluation Scale (Marin et al., 1991) and Lille Apathy Rating Scale (Sockeel et al., 2006)). These forms of assessment rely on the individual to recognise changes in their own behaviour and have limited sensitivity to track changes in motivated behaviour over time. These subjective methods cannot be directly translated to rodent tasks, however some behavioural tasks that assess motivated behaviour in rodents have been successfully back translated to human study, including the progressive ratio (PR) (Chelonis et al., 2011; Richardson and Roberts, 1996) and effort for reward tasks (EfR) (Heath et al., 2015; Salamone et al., 1991; Treadway et al., 2009). The PR task requires the rodent to put in increasing amounts of effort for each subsequent reward, while EfR gives the rodent the choice between an easy to obtain but low value food reward or a harder to obtain but higher value reward.

Emotional blunting, defined as a diminished response to emotionally salient stimuli, is a core feature of apathy but is often overlooked in favour of effort-based paradigms, particularly in rodent models (Magnard et al., 2016). Methods commonly used to study anxiety or depression-related behaviours may provide an indication of relevant aspects of emotional behaviour in ageing although studies are limited. Some groups have reported changes in anxiety/depression in models of neurodegenerative disorders (Taylor et al., 2010) and more recently, healthy aging (Shoji and Miyakawa, 2019). One study used changes in fear processing as a model of emotional blunting in a rodent model of schizophrenia (Pietersen et al., 2007). Given the multi-domain nature of apathy, elucidating an apathy phenotype requires a multifactorial approach that considers multiple levels, including emotional and motivational aspects.

In this study we used a cohort of normal healthy aged mice and tested them over a period of 9 months using behavioural assays specifically designed to probe different aspects of emotional behaviour, motivation, reward and cognition. At the end of the behavioural studies, animals were exposed to an acute restraint stress and both baseline and stress-induced corticosterone and post-mortem c-Fos expression were used to investigate the effects of ageing on stress reactivity. Together, these approaches were designed to test the hypothesis that normal aged mice develop emotional blunting and deficits in motivated behaviours consistent with an apathy-like phenotype.

## 2. Methods

### 2.1 Subjects

A cohort of 12 aged male C57bl/6J mice (15 mo at experiment onset and 24 mo by end of experimentation, 31.6-38.9g at onset and 33.6-40.0g by end) supplied by Eli Lilly and a cohort of C57bl/6J supplier, strain and sex-matched controls (3 mo at experiment onset and 12 mo by end of experimentation, 21.8-27.6g at experiment onset and 30.5-32.4g by end) supplied by Charles River were used. Sample size was based on previous behavioural studies using both spontaneous behavioural assays such as the open field arena and operant methods (Gourley et al., 2016; Rex et al., 1998). However, this type of aging work is novel and effect sizes may be smaller than those more typically seen with manipulations in these assays. Prior to arrival in Bristol, aged mice were group-housed in enriched caging and fed a restricted diet of 3g to promote healthy aging. On arrival in Bristol, all mice were individually housed in standard housing with a plastic house, cardboard tube and wooden chew block. They were kept in temperature-controlled conditions (21°C) and a 12:12 hour light-dark cycle (lights OFF at 0815, lights ON at 20:15). Standard lab chow (Purina, UK) was provided *ad libitum* apart from during operant training where diet was restricted diet of 2g chow per mouse. Weights were monitored at least once a week and maintained to at least 85% of their free feeding weight relative to their normal growth curve. Water was provided *ad libitum.* The same cohorts of animals were used for all studies and a timeline indicating the order of testing is given in **fig.1**. As the tests were performed sequentially there is the potential that increasing age may have affected some measures and hence the design randomised the tasks between measures of motivation-related and emotional behaviour. It should be noted that this design does not fully mitigate an effect of order of testing or increasing age of the animals. Within each behavioural assay, age was counterbalanced to account for time of day differences. Where possible, the experimenter was blind to age group though this was not always possible due to obvious physical differences between age groups. All experiments took place in the animals’ active phase and were performed in accordance with the Animals (Scientific Procedures) Act (UK) 1986 and were approved by the University of Bristol AWERB.

**Figure. 1.**
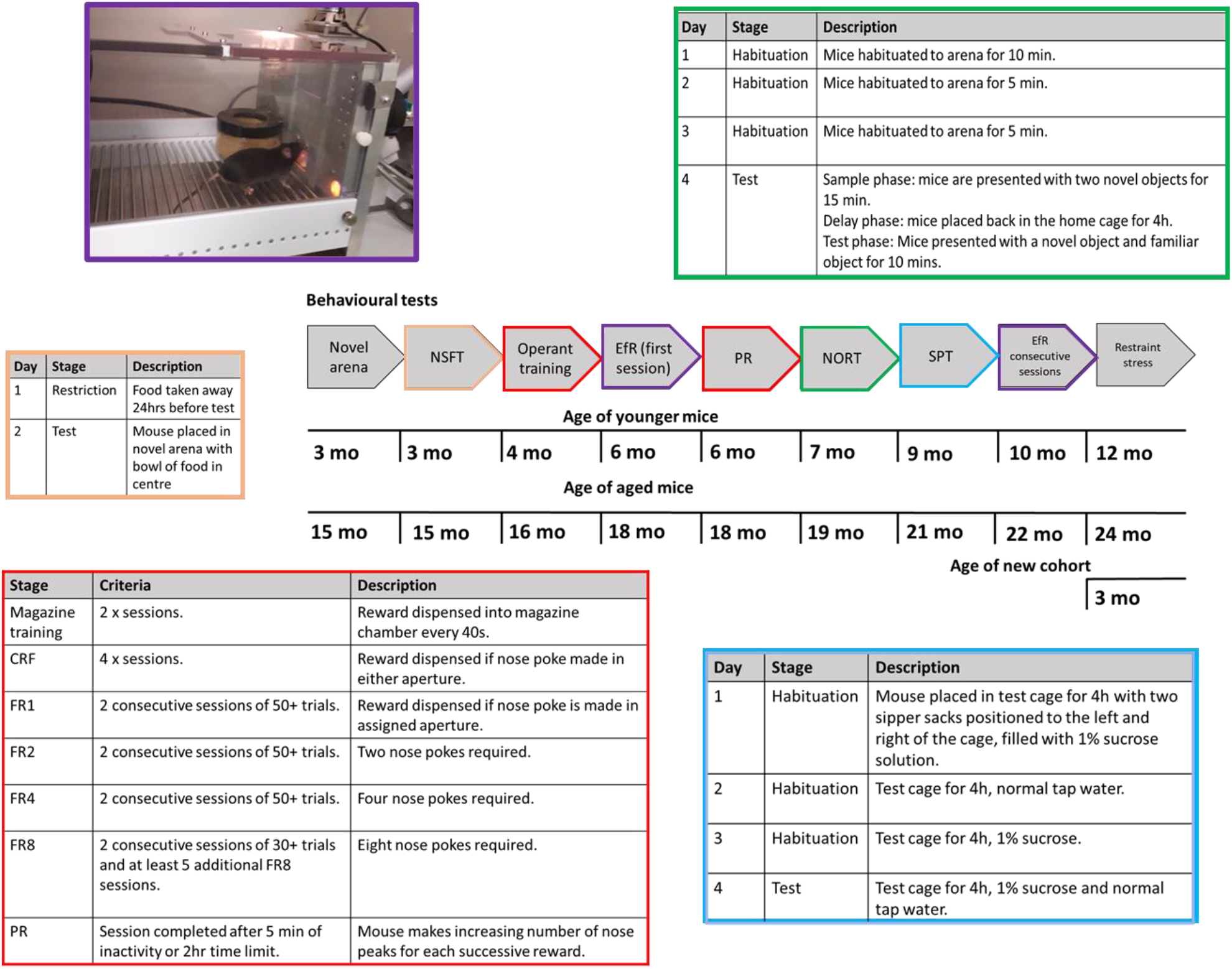
Experimental timeline and training protocols. Experimental timeline illustrating the different behavioural procedures used and when these occurred. Where more complex behavioural tasks have been used, additional information about training is also included and colour coded to link to the specific task within the timeline. Abbreviations: CRF-continuous reinforcement, EfR-effort for reward, FR-fixed ratio, NORT-novel object recognition test, NSFT-novelty supressed feeding test, PR-progressive ratio, SPT-sucrose preference test.

### 2.2 Exploration of a novel arena

Mice were placed in a novel, circular open field arena, 85 cm in diameter. Movement was captured for 15 min using a Logitech HD Pro Webcam c920 suspended 1 m above the arena. Videos were analysed using Ethovision Xt10 software (Noldus Information Technology, Wageningen) and total distance travelled (cm) and velocity (cm/s) were output.

### 2.3 Novelty suppressed feeding test

The protocol used was similar to that reported by (Shephard and Broadhurst, 1982). Mice were food deprived for 24 h before being placed in the left-hand corner of a novel arena (clear Perspex, 40 cm^2^ and lined with sawdust). A ceramic bowl of food was placed in the centre of the arena. Time taken for the mice to approach the bowl and to eat were manually recorded. The sawdust was shaken, the food bowl was cleaned, and fresh food added between mice.

### 2.4 Operant training

Training was similar to that reported by (Stuart et al., 2019). Mice were trained in sound-proof operant boxes (Med Associates Inc) which were run on Klimbic software (Conclusive Solutions Ltd., UK). Each operant box consisted of two nose poke apertures positioned either side of a centrally located food magazine. The magazine was connected to a reward pellet dispenser (20 mg rodent tablet, TestDiet, Sandown). Mice were run once per day during their active phase (9:00-17:00). Mice first learned to associate the magazine with the delivery of a reward pellet (one pellet every 40 sec) over two 30 min sessions. Mice then progressed to continuous reinforcement training (CRF) where a response made in either the left or right assigned nose poke aperture resulted in a single reward pellet. The magazine was illuminated until the mouse collected the pellet. In the third stage of training, the mice were required to advance through ascending fixed ratios (FR) of reinforcement making responses in either the left or right aperture only (counterbalanced across cohorts). The mice progressed through FR1, 2, 4 and 8, where the number refers to the number of nose pokes into the active aperture required for the delivery of one reward pellet. Mice completed each FR level when they obtained 50+ pellets over two consecutive sessions until FR8 where criteria was 30+ pellets over two consecutive sessions. Once all mice were trained, mice then completed a minimum of 5 additional FR8 sessions to manage differences in time to train (**fig.1**).

### 2.5 Effort for reward task

Directly after FR training was completed mice underwent a single FR8 session with access to low value, *ad libitum* powdered standard lab chow presented in a pot, placed in front of the inactive nose poke aperture, similar to that previously described by (Salamone et al., 1991). The pot was accessed via a ½ inch hole in the lid. Chow consumed was measured using change in weight of chow pre and post session, in g. Later in the behavioural battery (**fig.1**) mice were re-tested in the EfR task but this time over 5 consecutive sessions.

### 2.6 Progressive ratio task

Mice were tested in a progressive ratio (PR) task, in which each successive reward (*n*) required an increasing number of nose pokes using the algorithm F(*n*)=5 x EXP(0.2*n*)-5 (Roberts and Richardson, 1992). A PR session consisted of a maximum of 100 trials or 120 minutes and included a 1 second intertrial interval. 5 mins of inactivity ended the session. This test was conducted under both *ad libitum* and food restricted conditions. Breakpoint was defined as the last ratio competed before 5 min period of inactivity. Mice underwent one session of the PR task under each feeding condition. A PR session was preceded by an FR8 session to check for stability in performance. Preceding the described PR conditions mice were tested with a PR session that lasted a maximum of 60 mins or 10 mins of inactivity. However, it was clear a breakpoint would not be reached under these conditions (data not shown).

### 2.7 Consumption test

Mice were food restricted overnight and the following day were presented with free access to either powdered chow or reward pellets for 10 min over two different days in the home cage. Total amount consumed in g was calculated.

### 2.8 Novel object recognition test

This protocol was similar to that previously developed by (Ennaceur and Delacour, 1988). Mice were habituated to a Perspex arena (40 cm^2^) lined with paper liner for 10 min (day 1) or 5 min (day 2 and 3). On day 4, animals were tested using a sample phase where each mouse was presented with two novel objects for 15 mins. Mice were returned to their home cage during the 4h delay phase before being returned to the same arena for the test phase. Each animal was presented with both a novel and familiar object for 10 mins (**fig.1**). During both the sample and test phase, exploration was captured using a Logitech c920 webcam and then scored manually using DOSBox 0.74 software. Criteria for inclusion in the analysis was 20+ secs of total exploration in the sample phase. A discrimination ratio was calculated using *(time spent exploring novel object – time spent exploring familiar object)/time spent exploring novel object.*

### 2.9 Sucrose preference test

The protocol used was similar to that of (Willner et al., 1987). Mice were water restricted overnight for ~16h before both the habituation and test sessions. On day 1 and 3 mice were placed into test cages which contained sawdust and a cardboard tube and were presented with a 1 % sucrose solution in two sipper sacks (Edstrom-Avidity Science) with drip-free drinking valves, placed to the left and the right of the cage. On day 2 mice were presented with two sipper sacks containing normal tap water. On the test day they were presented with one sipper sack containing 1 % sucrose solution and the other containing tap water (**fig.1**). Position was counterbalanced across the cohort, and was swapped at 1h, and 2h. The habituation and test sessions lasted 4h and liquid consumed was weighed at the 1h, 2h and 4h time points. Sucrose preference was calculated using *(total amount of sucrose consumed/ (total sucrose + total water consumed*)) × 100.

### 2.10 Acute restraint stress

At the end of behavioural experiments, groups were approximately 12 months and 24 months old. As such an additional cohort was brought in as a young group. N = 12 male C57bl/6J mice from Charles River (3 mo at experiment onset) were singly housed in the same conditions as above. The new mice weighed 23.2-26.9g at experiment end. One mouse from the oldest group died before this final experiment.

Mice were restrained in a restraint tube (Ad Instruments Ltd) and their tail warmed for 3 minutes on a heat pad to promote blood flow in the tail vein. The tail vein was then opened with a 25G needle (Sigma-Aldrich, Germany) and blood was collected using a Mitra Microsampling sponge (Neoteryx, USA), with a calculated average blood wicking volume of 10 μl. The mouse was left in the restraint tube for a further 24 mins, before having the tail vein warmed again for 3 mins and another blood sample was taken. As an additional measure faecal pellets during testing were counted. Sampling took place between zeitgeber time (ZT) 17-20. Blood samples were stored at room temperature with a bag of desiccant until analysis.

### 2.11 Mass spectrometry analysis of corticosterone

Following sample extraction corticosterone levels were analysed using high-performance liquid chromatography/electrospray ionization tandem mass spectrometry (HPLC-ES/MS-MS) (for full experimental details see supplementary methods).

### 2.12 Tissue collection

Mice were returned to their home cage after the 30 min restraint stress and then killed by cervical dislocation after a further 60 min (90 min after onset of restraint stress). Brains were immediately removed and placed in 4 % paraformaldehyde (PFA) solution prepared in phosphate buffer saline (PBS) overnight and then transferred to a 25 % sucrose solution (Sigma Aldrich, UK) prepared in PBS. They were then frozen in OCT (Cryomatrix, Thermofisher) and stored at –20 °c.

### 2.13 c-Fos immunohistochemistry

Brains were sliced in 40 μm coronal sections using a freezing microtome (Reichert, Austria). Sections blocked for 30 mins in TBS-T (Trizma buffer saline with 0.1 % Triton X) with 3 % normal goat serum (Vector Laboratories). Sections were incubated with primary antibody rabbit anti c-Fos (1:4000, ABE457, Merck Millipore) overnight at 4° c. Sections were then incubated with secondary antibody goat anti-rabbit (1:500, Alexa Fluor 488, Invitrogen) for 3 hours and then with 4’,6-diamidino-2-phenylindole (DAPI) for a further 3 minutes (**for full experimental details see supplementary methods**).

Images were taken on a Leica widefield microscope with DFC365 FX camera with LasX software. The paraventricular nucleus of the hypothalamus (PVN) was captured with a 10x objective and the central (CeA) and basolateral amygdala (BLA) with a 5x objective. A picture of the PVN was taken across 3 different sections, bregma level −0.58 to −0.94 mm. A picture of the amygdala (left or right) was taken across 3 different sections, bregma level (−1.22 to −1.40 mm). Some brains/sections were lost due to tissue damage. N = 10 per group amygdala was obtained. N = 10 PVN for aged and young group were obtained, N = 9 for middle aged group. c-Fos positive cells that co-localised with DAPI were counted manually using ImageJ cell counter. Contrast was adjusted uniformly across all images. c-Fos count was normalised by dividing c-Fos count by area of region.

### 2.14 Data analysis

Repeated measures (RM) two-way ANOVA, one-way ANOVA, independent samples t-test or the Kruskal-Wallis test (for non-parametric faecal count data) were performed where appropriate. Where significant main effects or interactions were observed (p < 0.05) these are reported in the results section with appropriate post-hoc pairwise comparisons to explore group differences. Where main effect or interaction statistics found a trend level effect (p < 0.1) these are mentioned in the text but were not further analysed. When conducting ANOVA tests, where data did not meet the assumption of sphericity, the Huynh-Feldt correction was used to adjust degrees of freedom. When conducting single independent t-tests, Levene’s Test for Equality of Variances was used to determine whether equal variances were assumed with correction being applied if this was not valid. Analysis was performed on IBM SPSS statistics v.24 for Windows and all graphs were made using GraphPad Prism 8.3.0 for Windows.

## 3. Results

### 3.1 Aged mice show reduced exploration of a novel arena but show no deficit in the novel object recognition task

In a novel arena, aged mice covered less distance and had a slower mean velocity than younger mice (t_(22)_ = 2.536, p = 0.019 and t_(22)_ = 2.613, p = 0.016 respectively, independent t-test) **(fig.2A&B)**. When data was split into 5 min time bins there was a main effect of time (F_(2,44)_ = 35.330, p = 7.06 x 10^-10^, RM two-way ANOVA) where distance travelled reduced with time. There was also a main effect of age group (F_(1,22)_ = 6.434, p = 0.019) but there was no group*time interaction (p > 0.05). Pairwise comparisons revealed no difference in distance travelled between groups in the first 5 mins, but a difference emerged in the second- and third-time bin p = 0.013 and p = 0.049 respectively **(fig.2C)**.

**Figure. 2.**
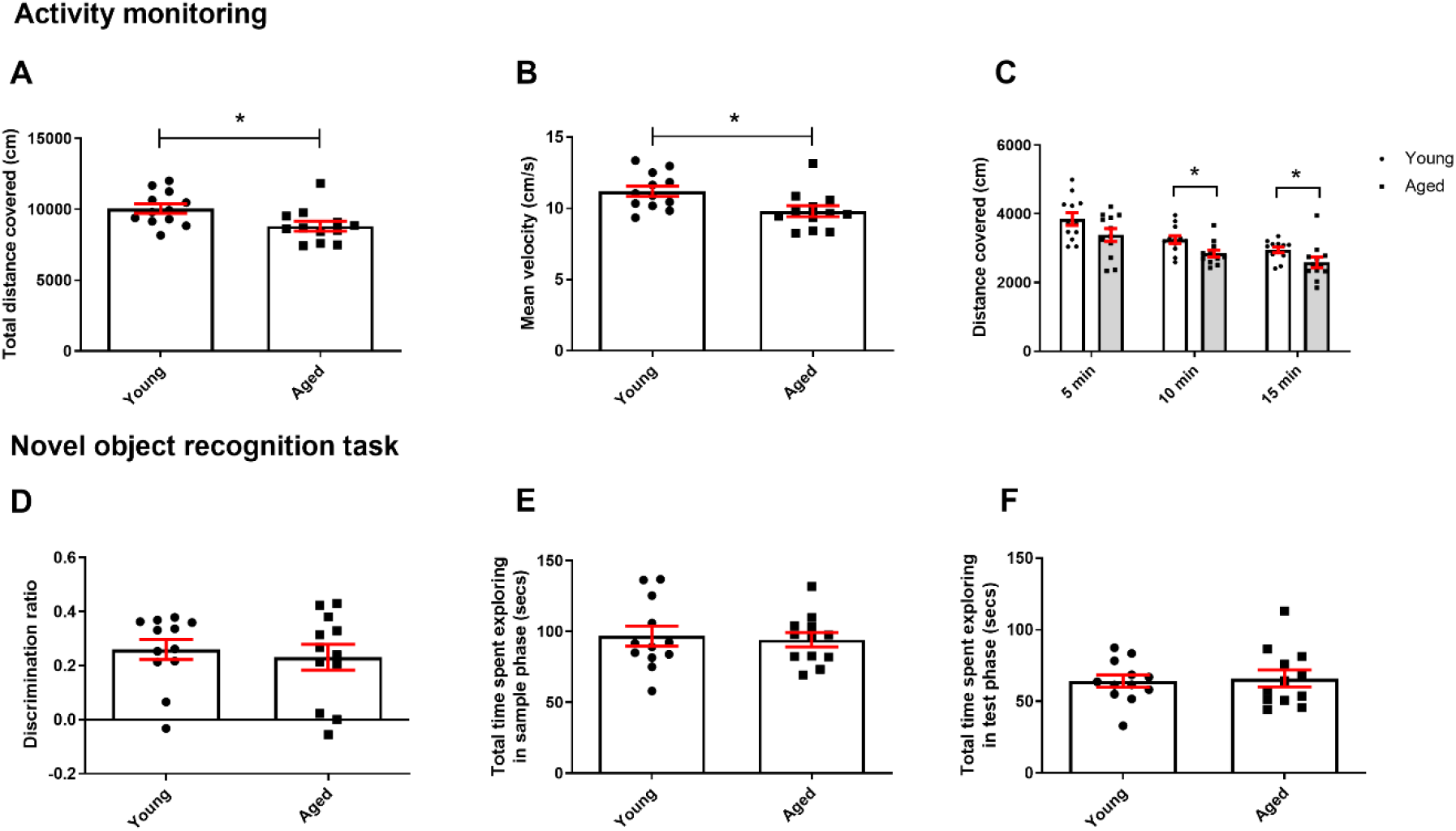
Aged mice show reduced exploration in a novel arena but normal cognition in the novel object recognition test. **A** Aged mice covered less distance overall in a novel environment compared to younger mice (independent t-test). **B** Aged mice were slower than younger mice (independent t-test). **C** Distance covered was not different between groups in the first 5 mins of the task but was by 10 and 15 mins. Mice also covered less distance over time (RM two-way ANOVA with pairwise comparisons). **D** There was no difference in discrimination ratio during the test phase of the NOR task. Bars are mean ± SEM, with data points overlaid. **E** There was no difference in time spent exploring objects in the sample phase of the NORT between groups, **F** or in the test phase. *p < 0.05, ###p < 0.001-refers to main effect of time. N = 12 per group.

In the NOR task, there no difference in discrimination ratio between groups (p > 0.05) **(fig.2D)** and there was no difference in exploration of the objects between groups during the sample phase **(fig.2E)** or test phase **(fig.2F)**.

### 3.2 Aged mice show changes in motivated behaviour

Under food restriction aged mice ended the PR session at a lower final ratio completed (t_(12.464)_ = 4.563, p = 0.001, independent t-test), however mice failed to reach a true breakpoint i.e. they kept working until session time limit. Analysis of time taken to complete each ratio within the PR session showed a main effect of ratio (F_(1.403, 30.865)_ = 22.812, p < 0.0001, RM two-way ANOVA), where time taken to complete a ratio increased with ratio. There was also a ratio*age group interaction (F_(1.403, 30.865)_ = 13.581, p < 0.0001) and a main effect of age (F_(1,22)_ = 19.960, p < 0.0001). Ratio 62 was the final ratio all mice completed so was used as a cut off for analysis. Pairwise comparisons showed younger mice completed ratio 6-62 faster than aged mice (p ≤ 0.016) **(fig.3A)**. Under *ad libitum* feeding conditions 8 young mice and 9 aged mice reached breakpoint i.e. ended session with 5 mins of inactivity. There was no difference in breakpoint between groups (p > 0.05). However when all mice were added to the analysis, a difference emerged (t_(16.108)_ = 2.619, p = 0.019). There was no difference in time taken to complete each ratio between groups (up to ratio 40). RM ANOVA was not possible for this analysis due to variation in final ratio completed so a series of independent t-tests were conducted instead (t_(15)_ ≤ 1.565, p ≥ 0.113, independent t-tests) **(fig.3B)**.

**Figure. 3.**
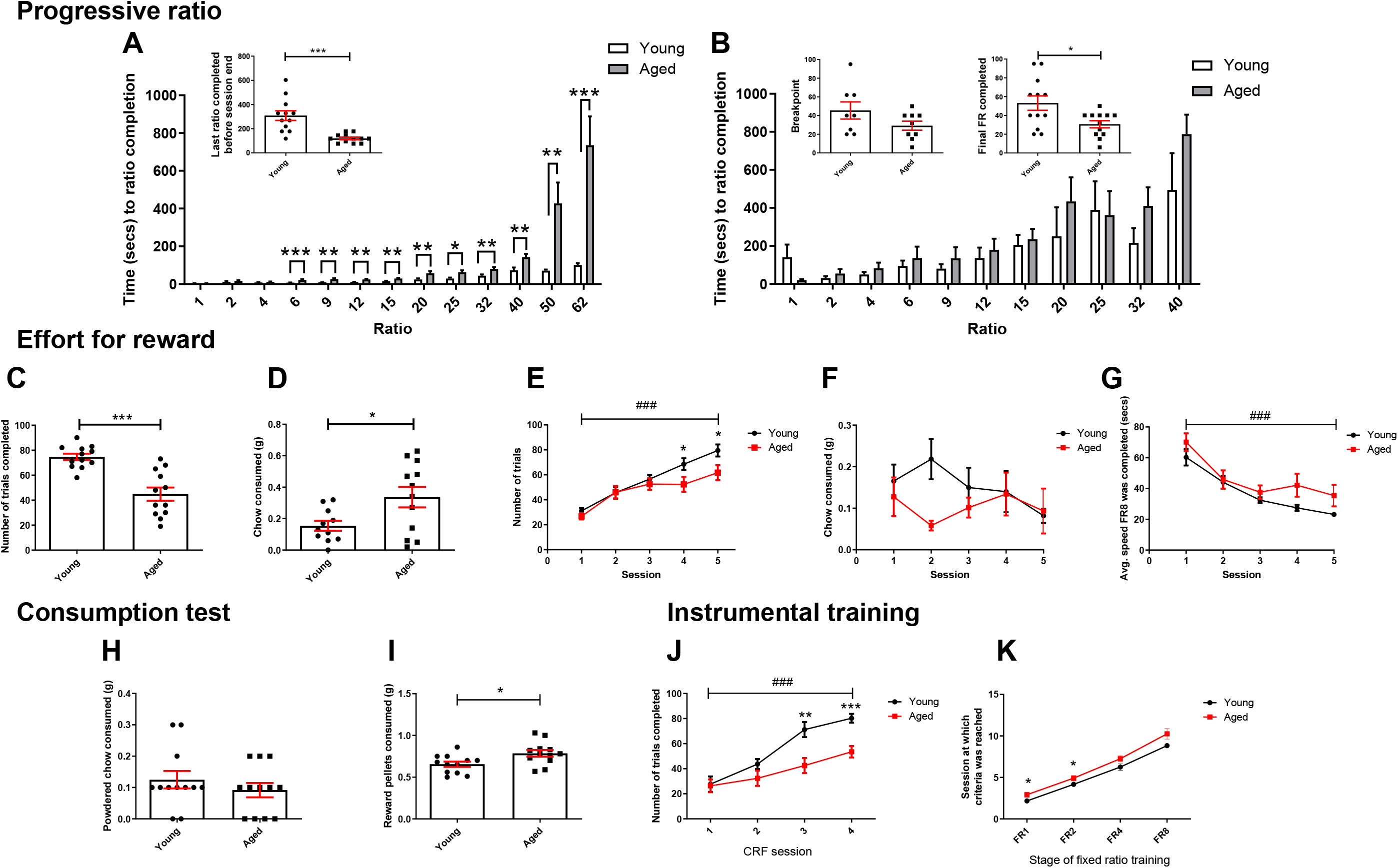
Aged mice show changes in motivation under different conditions. **A** Under food restriction, aged mice completed the task on a lower final ratio in the progressive ratio schedule than younger mice (independent t-test). Younger mice completed ratios 6-62 faster than aged mice (RM two-way ANOVA with pairwise comparisons). **B** Under *ad libitum* feeding conditions and for animals which achieved a true breakpoint, there was no difference between age groups (independent t-test). However, when all mice were included in the analysis, aged mice finished on a lower ratio than younger mice (independent t-test). There was no difference in speed across groups under free feeding conditions (independent t-tests). **C** on first exposure to the EfR task, aged mice completed less trials than younger mice (independent t-test). **D** In the same session, aged mice consumed more chow (independent t test). **E** When the task was repeated over 5 consecutive days and at the end of the PR training and testing, aged mice completed less trials than younger mice only in sessions 4 and 5. Number of trials completed increased across the week in both groups (p < 0.05, p < 0.001, RM two-way ANOVA with pairwise comparison). **F** There was no effect of session or group on chow consumption. **G** Mice became faster at completing trials over sessions (p < 0.001, RM twoway ANOVA). **H** In a 10 min consumption test, there was no difference in chow consumed between groups (independent t-test). **I** Aged mice consumed more reward pellets than younger mice (independent t-test). **J** Aged mice completed less trials in CRF training session 4&5 (RM two-way ANOVA with pairwise comparison). **K** Aged mice took more sessions to complete FR1 and FR2 stages than younger mice (RM two-way ANOVA with pairwise comparisons). Bars are mean ± SEM with data points overlaid.* p<0.05, **p<0.01, ***p<0.001, ###p<0.001-refers to main effect of session. N = 10-12 per group.

On first exposure to the effort for reward task, aged mice obtained less of the high value reward pellets than younger mice (t_(15.982)_ = 5.088, p = 0.0001, independent t-test) but consumed more of the low value chow (t_(15.819)_ = 2.453, p = 0.023). One younger mouse was excluded for excessive digging in chow **(fig.3C&D)**. In a subsequent 5 day test, analysis of number of trials completed over sessions showed a main effect of session (F_(3.22, 70.833)_= 41.553, p < 0.0001, RM two-way ANOVA) and there was a session*group interaction (F_(3.22, 70.833)_ = 2.73, p = 0.047). There was a trend level effect of age (F_(1,22)_ = 3.038, p = 0.095). Post-hoc pairwise comparison showed younger mice completed more trials only in the final two sessions (p = 0.046 and p = 0.032 respectively) **(fig.3E).** However, there was no effect of session or group on consumption of chow (p > 0.05) although there was a trend towards a chow*group interaction (F_(3.59, 75.4)_ = 2.237, p= 0.08) **(fig.3F)**. n= 1 young mouse was excluded for digging in chow over consecutive sessions. In a single instance of digging, value was replaced with group mean to permit ANOVA analysis (n = 1 young mouse). Analysis of average speed (speed at which FR8 was completed, averaged over a session) across the sessions showed a main effect of session on speed, where speed increased over sessions in both groups (F_(2.302, 50.636)_ = 34.476, p < 0.0001). However, there was no age*session interaction or effect of age (p > 0.05) (**Fig.3G**). Consumption tests showed aged mice consumed more reward pellets in 10 mins than younger mice (t_(22)_ = 2.587, p = 0.0168, independent t-test) while consumption of chow did not differ (p > 0.05) **(Fig.3H&I)**.

Analysis of CRF training performance revealed a main effect of session on number of trials completed (F_(3,66)_ = 44.471, p <0.0001, RM two-way ANOVA), as well as a session*age group interaction (F_(3,66)_ = 5.815, p = 0.0014) and a main effect of age (F_(1,22)_ = 8.365, p = 0.008). There was no difference in performance between groups in the first two sessions, but younger mice completed more trials in the final two sessions (p ≤ 0.003). **(fig.3J)**. Analysis of FR training performance showed a main effect of FR stage on number of sessions to criteria (F_(3,66)_ = 374.63, p < 0.0001). There was also a main effect of age on number of sessions to FR stage completion (F_(1,22)_ = 5.539, p = 0.028) but there was no FR stage*age interaction (p > 0.05). Aged mice took more sessions to complete FR1 and FR2 than younger mice, p = 0.041) but not FR4 and FR8 p = 0.078 and p = 0.058 respectively, though there was a trend **(fig.3K)**.

### 3.3 Aged mice show changes in hedonic behaviour, anxiety-like behaviour and stress reactivity

Analysis of % sucrose preference showed a main effect of time (F_(1.781, 39.191)_ = 12.146, p < 0.0001, RM two-way ANOVA) where sucrose preference increased with time. There was also a time*group interaction (F_(1.781, 39.191)_= 7.076, p = 0.003) and an effect of age group (F_(1,22)_ = 16.692, p < 0.0001) **(fig.4A)**. There were 3 leaks in water sacks during testing (2 aged, 1 young) and where this occurred the value for this sack was replaced with the group mean to permit RM ANOVA analysis. Post hoc pairwise comparison showed younger mice had a higher sucrose preference than aged mice at each time point (p ≤ 0.043).

**Figure. 4.**
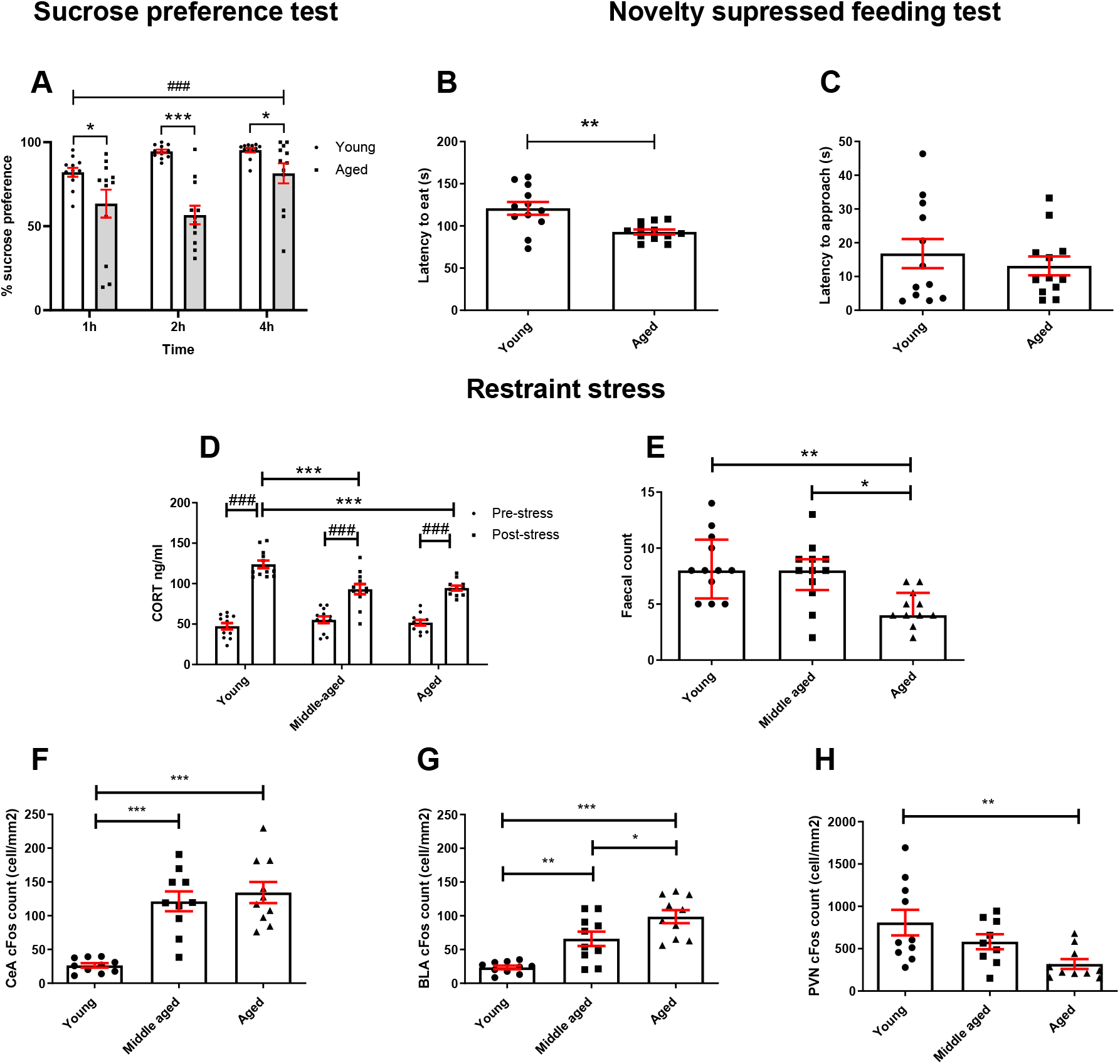
Aged mice show changes in reward sensitivity and stress reactivity. **A** Aged mice showed a reduced sucrose preference compared to younger mice across all time points, and preference changed across time (RM two-way ANOVA with pairwise comparisons). **B** In the NSFT, aged mice took less time to eat from the bowl than younger mice (independent t-test). **C** There was no difference in latency to approach bowl between groups (independent t-test). **D** Following 30 min restraint stress, CORT was increased in all groups but aged mice showed a reduced CORT response to stress compared to younger mice but not middle aged (RM two-way ANOVA with pairwise comparisons). **E** Aged mice had a lower faecal count following stress compared to young and middleaged groups (Kruskal-Wallis test with Dunn’s post-hoc (bars are median with interquartile range). **F** Following stress, middle-aged and aged mice had a greater c-Fos count in the CeA compared to younger mice (p<0.001, one-way ANOVA with Tukey’s post-hoc analysis). **G** Aged mice had a greater BLA c-Fos count compared to middle aged and young mice. Younger mice had a lower c-Fos count in the BLA compared to middle aged mice (one-way ANOVA with Tukey’s post-hoc analysis). **H** Younger mice had a greater c-Fos count in the PVN compared to aged mice (one-way ANOVA with Tukey’s post-hoc analysis). Unless otherwise indicated bars are mean ± SEM with data points overlaid. *p<0.05, **p<0.01, ***p<0.001, ###p<0.001-refers to main effect of time or pairwise effect of stress. N = 9-12 per group.

Aged mice took less time to eat from the bowl in the novelty supressed feeding test (t_(4.208)_ = 3.411, p = 0.004, independent t-test) **(Fig.4B)** but latency to approach the bowl was not different (p > 0.05) **(Fig.4C)**. Under acute restraint stress, baseline corticosterone was similar for all groups but aged animals had a lower corticosterone response to restraint (main effect of stress F_(1,32)_ = 179.608, p < 0.0001, age F_(1,32)_ = 5.772, p = 0.007 and stress*group interaction F_(2,32)_ = 9.862, p = 0.0005, RM twoway ANOVA). Pairwise comparison revealed CORT was increased following stress in all groups, but this was higher in the young mice (young vs aged or middle-aged p = 0.001 and p = 0.0003 respectively) **(fig.4D)**. There was also an effect of age on faecal count (H_(2)_ = 12.424, p = 0.002, Kruskal-Wallis test). Dunn’s post-hoc test showed aged mice had lower faecal count than middleaged mice (p = 0.011) and young mice (p = 0.004) **(fig.4E)**. Post-mortem analysis of neuronal activation following restraint stress showed CeA c-Fos count was higher in both the aged and middle-aged animal (main effect of age F_(2,29)_ = 22.102, p < 0.0001, one-way ANOVA) with Tukey’s post hoc analysis (p < 0.0001 vs younger mice) **(fig.4F)**. There was also an effect of age on BLA c-Fos count (F_(2,29)_ = 19.915, p < 0.0001), where middle-aged and young mice had a lower BLA cFos count than aged mice (p = 0.029 and p < 0.0001 respectively). Middle-aged mice also had a higher BLA cFos count than younger mice (p = 0.004) **(fig.4G)**. PVN c-Fos count was higher in younger mice compared to aged mice (main effect of age F_(2,28)_ = 5.326, p = 0.012, Tukey’s post hoc, p = 0.008) **(fig.4H)**.

## 4. Discussion

We observed that aged mice displayed behavioural deficits in both motivation and affect which suggests they may develop a phenotype with at least some characteristics of human apathy. Using measures related to motivation, we observed that aged mice were less active in a novel environment, responded with reduced vigour in the progressive ratio task and were more likely to seek low value low effort reward under certain conditions. These effects did not appear to be related to changes in appetite, cognition, or gross changes in locomotor function. In the PR task, we noted that most animals did not reach a true break point, possibly due to higher levels of motivation resulting from food restriction. When we reduced motivation across all groups using free feeding, we found that the deficit in the aged animals related to mainly to rate of responding. Using a series of different measures of stress reactivity and affective behaviour we also found evidence of a blunted emotional response in aged mice. It is interesting to observe that the effects were not typical of that seen in models of depression (Planchez et al., 2019). In assays used to measure anxiety and depression-related behaviours we found effects suggesting lower levels of anxiety but reduced reward sensitivity. When we consider these alongside the blunted corticosterone and PVN c-Fos activation in response to restraint stress, we propose that aged mice also show emotional blunting. The following discussion considers these two domains of apathy and the potential to use aged mice to better understand the symptom of apathy.

### 4.1 Aged mice show a reduction in goal-directed behaviour

Aged mice were slower and covered less distance in the open field arena. When the analysis was broken down by time bin, the effect was more apparent after 10 and 15 min suggesting that initial exploration of the environment did not differ although it should be noted the effects were marginal. These effects may be due to declining motivational drive to explore, though it is important to acknowledge that age-related deterioration in motoric capability may also play a role. Aging has been associated with psychomotor slowing (Tombaugh, 2004), and it has been suggested this slowing is due to a reduction in motivation and emotional arousal, which can be independent of age-related effects (Seidler et al., 2010).

During operant conditioning aged animals initially responded at a similar level but then failed to increase their rate of responding to the same level as the younger animals over time. Their rate of learning was parallel to the controls suggesting reduced vigour rather than a learning impairment. In the initial studies carried out under food restriction, aged animals appeared less motivated and completed less ratios for reward under a PR schedule of reinforcement. They were slower to complete the number of responses required for each ratio but a true breakpoint, in which the mouse gives up before the session ended was not reached. Many PR studies are limited by time rather than lack of responding (Heath et al., 2015) or a combination of both (Gourley et al., 2016). This may mean that conclusions are confounded by changes in rate of response and an assumption that the last ratio completed before the task times out is also break point. When the motivational state was reduced by providing food *ad libitum,* we found no difference in breakpoint or the speed at which each ratio was completed. This suggests rate of responding is driving the motivational deficit in aged mice and the effects are more apparent when control animals are in a high motivational state. It is common practise to use food restriction to motivate animals to learn operant tasks however this may create an artificial state which is more obvious in control animals thus biasing results for these types of task. These findings are in line with studies using a mouse model of Huntington’s disease, in which apathy is a core symptom, where they found a reduction in rate of response in a PR schedule of reinforcement, relative to controls (Heath et al., 2019; Oakeshott et al., 2012). This was also found in a study using Huntington’s disease patients with apathy, and apathy questionnaire scores negatively correlated with breakpoint scores (Heath et al., 2019).

An alternative way to look at motivation is the EfR task, an effort-based decision-making (EBDM) task which was run here using two different formats. The initial test was performed at the end of FR training and involved a single test session where aged mice completed less trials but consumed more chow, indicating preference for the lower effort option compared to younger mice. However, after animals had undergone multiple sessions under FR8 and PR schedules and, when the task was presented over consecutive days, a group difference emerged only in the final two sessions. There is a potential confound with this format which links to our previous PR data. Animals were run under food restriction, but mice are given food *ad libitum* over the weekend and food restricted across the week. In this second study, changes in chow consumption were not observed however, variability was much higher possibly resulting from excessive digging and waste from the chow bowl. Changes in EBDM have been reported in Schizophrenia patients and Parkinson’s disease patients with apathy (Chong et al., 2015; Le Heron et al., 2018b). However, much of pre-clinical work on this type of EBDM focusses on investigating the role of the dopaminergic system, rather than probing for apathy behaviour in disease/aging models (Le Heron et al., 2018a).

Together, these data suggest aged mice have a deficit in the activational phase of motivated behaviour, which is characterised by speed, vigour and persistence (Salamone et al., 2016). These changes appear not to be explained by changes in appetite or a cognitive impairment as no differences were observed in consumption tests or in the NOR task. There is conflicting evidence in the human literature whether motoric vigour is reduced in apathy, where some studies suggest vigour in pursuit of reward may be conserved in apathy patients, while other work points to a reduction (Le Heron et al., 2018a).

### 4.2 Aged mice exhibit emotional blunting

In the novelty supressed feeding test, aged mice were quicker to consume food in a novel environment suggested lower novelty-induced hyponeophagia. These findings are consistent with reduced anxiety and opposite to the effects seen in rodent models of depression where increased feeding latencies are consistently observed (Planchez et al., 2019). In contrast, aged animals had reduced reward sensitivity in the SPT. These findings are similar to the reward deficits seen in rodent models of depression (Willner, 2016) and are considered to reflect a measure of anhedonia. Studies have shown that elderly people have a reduction in reward sensitivity (Muhammed et al., 2016). A probabilistic reinforcement learning task showed that monetary loss had a larger impact on subsequent behaviour than gain, in elderly participants compared to younger participants (Hämmerer et al., 2010). This disruption may be due to age-related changes to the efficiency of dopaminergic and serotonergic neuromodulation, which has been demonstrated in fMRI studies based on learning and memory-related tasks (Eppinger et al., 2011; Schott et al., 2007). These behavioural findings provide an interesting contrast to depression-like phenotypes and suggest an emotional blunting rather than negative affective state.

To explore the apparent reduction in stress-reactivity seen in the NSF test we also tested animals using an acute restraint stress which reliably increases the stress hormone, corticosterone in rodents (Harizi et al., 2007; Nohara et al., 2016). There were no differences in corticosterone under baseline conditions and all groups showed an increase in plasma levels following 30 min restraint. Both groups of aged mice (12 and 24 mo) had a blunted CORT response and the older group also had a reduced faecal count. We also found age-related changes in the response to stress centrally, where c-Fos activation in the paraventricular nucleus of the hypothalamus (PVN) was reduced in the 24mo group compared to the youngest group, yet central (CeA) and basolateral amygdala (BLA) activation was increased in the aged groups. The PVN is a core part of the stress response, it’s activation by the limbic circuit and brainstem pathways activates the adrenocortical axis and results in the release of CORT (Benarroch, 2005). As such, this reduced activation could explain the reduced CORT response in the aged mice. The CeA plays a key role in fear, anxiety and the stress response, integrating behaviour and autonomic response to aversive stimuli (Day et al., 2005). Previous research has shown that there is relatively little activation of this region in response to processive stressors, including restraint stress and is more responsive to homeostatic disruption or systemic stress (Day et al., 2005). However, aged mice show a greater activation of the BLA/CeA during stress compared to the young mice. The reason for this and its functional significance is unclear. However, it has been previously suggested that changes to the functional gating of limbic information by local PVN projections may explain age-related changes to the HPA axis (Herman et al., 2002). Stress reactivity and active coping in response to aversive experiences has previously been shown to reduce with age in mice (Oh et al., 2018) and extend beyond mice to rat and human studies (Brugnera et al., 2017; Buechel et al., 2014). There is some variability in findings possibly due to a lack of standardisation across studies including the nature and intensity of the stressor (Novais et al., 2017; Segar et al., 2009).

## 5. Conclusion

By using a battery of behavioural tests that probe aspects of motivated and emotional behaviour, we show that aged mice have deficits in behaviours relating to multiple domains of apathy. Specifically, deficits in goal-directed behaviour measured by the exploration of a novel arena, PR task and EfR task map onto the auto-activation/behavioural component of apathy. Crucially, the observed behavioural changes were not explained by general appetite differences, cognitive impairments or overt changes in motor function. A reduced stress response, evidenced by behavioural analysis, blunted stress-induced CORT and reduced PVN activation, and reduced reward sensitivity provide strong evidence for emotional blunting in aged mice. Together, these data suggest naturally aged mice have the potential to provide a model to investigate the underlying neurobiology of apathy. The findings relating to stress reactivity add to previously published work (Oh et al., 2018) suggesting an age-related decline in active coping including the hormonal response to acute stress. This raises an important issue when considering data obtained from stress-driven cognitive tasks such as the commonly used Morris Water Maze (Morris, 1981). As this task is motivated by the aversive nature and stress-inducing effects of the animal being placed into water, changes in stress reactivity could influence learning independent of a specific cognitive impairment.

## Abbreviations

BLA: basolateral amygdala
CeA: central amygdala
CORT: corticosterone
EBDM: effort-based decision making
EfR: effort for reward
ES: Electrospray
HLPC: high performance liquid chromatography
MS: mass spectrometry
NORT: novel object recognition test
NSFT: novelty supressed feeding test
PR: progressive ratio
PVN: paraventricular nucleus of the hypothalamus
SPT: sucrose preference test

## Acknowledgements

We acknowledge the help of Julia Bartlett, Dr Claire Hales and Dandri Aly Purawijaya for assisting in aspects of behavioural testing/tissue preparation, Dr Sandra Sossick, Eli Lilly for conducting the mass spectrometry analysis of corticosterone and Dr Christopher Holton, Eli Lilly for the supply of aged mice. We acknowledge the Wolfson Bioimaging Facility for use of the brightfield microscope.

## References

Benarroch, E.E., 2005. Paraventricular nucleus, stress response, and cardiovascular disease. Clinical Autonomic Research 15(4), 254–263.

Brugnera, A., Zarbo, C., Adorni, R., Gatti, A., Compare, A., Sakatani, K., 2017. Age-Related Changes in Physiological Reactivity to a Stress Task: A Near-Infrared Spectroscopy Study, in: Halpern, H.J., LaManna, J.C., Harrison, D.K., Epel, B. (Eds.), Oxygen Transport to Tissue XXXIX. Springer International Publishing, Cham, pp. 155–161.

Buechel, H., Popovic, J., Staggs, K., Anderson, K., Thibault, O., Blalock, E., 2014. Aged rats are hypo-responsive to acute restraint: Implications for psychosocial stress in aging.

Chelonis, J.J., Gravelin, C.R., Paule, M.G., 2011. Assessing motivation in children using a progressive ratio task. Behavioural Processes 87(2), 203–209.

Chong, T.T.J., Bonnelle, V., Manohar, S., Veromann, K.-R., Muhammed, K., Tofaris, G.K., Hu, M., Husain, M., 2015. Dopamine enhances willingness to exert effort for reward in Parkinson’s disease. Cortex 69, 40–46.

Day, H.E.W., Nebel, S., Sasse, S., Campeau, S., 2005. Inhibition of the central extended amygdala by loud noise and restraint stress. The European journal of neuroscience 21(2), 441–454.

Ennaceur, A., Delacour, J., 1988. A new one-trial test for neurobiological studies of memory in rats. 1: Behavioral data. Behavioural Brain Research 31(1), 47–59.

Eppinger, B., Hämmerer, D., Li, S.-C., 2011. Neuromodulation of reward-based learning and decision making in human aging. Annals of the New York Academy of Sciences 1235, 1–17.

Gerritsen, D.L., Jongenelis, K., Steverink, N., Ooms, M.E., Ribbe, M.W., 2005. Down and drowsy? Do apathetic nursing home residents experience low quality of life? Aging & Mental Health 9(2), 135–141.

Gourley, S.L., Zimmermann, K.S., Allen, A.G., Taylor, J.R., 2016. The Medial Orbitofrontal Cortex Regulates Sensitivity to Outcome Value. The Journal of Neuroscience 36(16), 4600–4613.

Hämmerer, D., Li, S.-C., Müller, V., Lindenberger, U., 2010. Life Span Differences in Electrophysiological Correlates of Monitoring Gains and Losses during Probabilistic Reinforcement Learning. Journal of Cognitive Neuroscience 23(3), 579–592.

Harizi, H., Homo-Delarche, F., Amrani, A., Coulaud, J., Mormède, P., 2007. Marked genetic differences in the regulation of blood glucose under immune and restraint stress in mice reveals a wide range of corticosensitivity. Journal of Neuroimmunology 189(1), 59–68.

Heath, C.J., Bussey, T.J., Saksida, L.M., 2015. Motivational assessment of mice using the touchscreen operant testing system: effects of dopaminergic drugs. Psychopharmacology 232(21-22), 4043–4057.

Heath, C.J., O’Callaghan, C., Mason, S.L., Phillips, B.U., Saksida, L.M., Robbins, T.W., Barker, R.A., Bussey, T.J., Sahakian, B.J., 2019. A Touchscreen Motivation Assessment Evaluated in Huntington’s Disease Patients and R6/1 Model Mice. Frontiers in Neurology 10(858).

Herman, J.P., Tasker, J.G., Ziegler, D.R., Cullinan, W.E., 2002. Local circuit regulation of paraventricular nucleus stress integration: Glutamate–GABA connections. Pharmacology Biochemistry and Behavior 71(3), 457–468.

Ishii, S., Weintraub, N., Mervis, J.R., 2009. Apathy: A Common Psychiatric Syndrome in the Elderly. Journal of the American Medical Directors Association 10(6), 381–393.

Kamat, R., Woods, S.P., Marcotte, T.D., Ellis, R.J., Grant, I., and the, H.I.V.N.R.P.G., 2012. Implications of Apathy for Everyday Functioning Outcomes in Persons Living with HIV Infection†. Archives of Clinical Neuropsychology 27(5), 520–531.

Le Heron, C., Holroyd, C.B., Salamone, J., Husain, M., 2018a. Brain mechanisms underlying apathy. Journal of Neurology, Neurosurgery & amp;amp; Psychiatry, jnnp-2018-318265.

Le Heron, C., Plant, O., Manohar, S., Ang, Y.-S., Jackson, M., Lennox, G., Hu, M.T., Husain, M., 2018b. Distinct effects of apathy and dopamine on effort-based decision-making in Parkinson’s disease. Brain: a journal of neurology 141(5), 1455–1469.

Levy, R., Czernecki, V., 2006. Apathy and the basal ganglia. Journal of Neurology 253(7), vii54–vii61.

Levy, R., Dubois, B., 2006. Apathy and the Functional Anatomy of the Prefrontal Cortex–Basal Ganglia Circuits. Cerebral Cortex 16(7), 916–928.

Magnard, R., Vachez, Y., Carcenac, C., Krack, P., David, O., Savasta, M., Boulet, S., Carnicella, S., 2016. What can rodent models tell us about apathy and associated neuropsychiatric symptoms in Parkinson’s disease? Translational Psychiatry 6(3), e753.

Marin, R.S., Biedrzycki, R.C., Firinciogullari, S., 1991. Reliability and validity of the apathy evaluation scale. Psychiatry Research 38(2), 143–162.

Morris, R.G.M., 1981. Spatial localization does not require the presence of local cues. Learning and Motivation 12(2), 239–260.

Muhammed, K., Manohar, S., Ben Yehuda, M., Chong, T.T.J., Tofaris, G., Lennox, G., Bogdanovic, M., Hu, M., Husain, M., 2016. Reward sensitivity deficits modulated by dopamine are associated with apathy in Parkinson’s disease. Brain: a journal of neurology 139(Pt 10), 2706–2721.

Nohara, M., Tohei, A., Sato, T., Amao, H., 2016. Evaluation of response to restraint stress by salivary corticosterone levels in adult male mice. J Vet Med Sci 78(5), 775–780.

Novais, A., Monteiro, S., Roque, S., Correia-Neves, M., Sousa, N., 2017. How age, sex and genotype shape the stress response. Neurobiology of Stress 6, 44–56.

Oakeshott, S., Port, R., Cummins-Sutphen, J., Berger, J., Watson-Johnson, J., Ramboz, S., Paterson, N., Kwak, S., Howland, D., Brunner, D., 2012. A mixed fixed ratio/progressive ratio procedure reveals an apathy phenotype in the BAC HD and the z_Q175 KI mouse models of Huntington’s disease. PLoS Curr 4, e4f972cffe982c970–e974f972cffe982c970.

Oh, H.-J., Song, M., Kim, Y.K., Bae, J.R., Cha, S.-Y., Bae, J.Y., Kim, Y., You, M., Lee, Y., Shim, J., Maeng, S., 2018. Age-Related Decrease in Stress Responsiveness and Proactive Coping in Male Mice. Front Aging Neurosci 10, 128–128.

Pietersen, C.Y., Bosker, F.J., Doorduin, J., Jongsma, M.E., Postema, F., Haas, J.V., Johnson, M.P., Koch, T., Vladusich, T., den Boer, J.A., 2007. An animal model of emotional blunting in schizophrenia. PLoS One 2(12), e1360–e1360.

Planchez, B., Surget, A., Belzung, C., 2019. Animal models of major depression: drawbacks and challenges. Journal of Neural Transmission 126(11), 1383–1408.

Rex, A., Voigt, J.P., Voits, M., Fink, H., 1998. Pharmacological Evaluation of a Modified Open-Field Test Sensitive to Anxiolytic Drugs. Pharmacology Biochemistry and Behavior 59(3), 677–683.

Richardson, N.R., Roberts, D.C.S., 1996. Progressive ratio schedules in drug self-administration studies in rats: a method to evaluate reinforcing efficacy. Journal of Neuroscience Methods 66(1), 1–11.

Roberts, D.C.S., Richardson, N.R., 1992. Self-Administration of Psychomotor Stimulants Using Progressive Ratio Schedules of Reinforcement, in: Boulton, A.A., Baker, G.B., Wu, P.H. (Eds.), Animal Models of Drug Addiction. Humana Press, Totowa, NJ, pp. 233–269.

Salamone, J.D., Steinpreis, R.E., McCullough, L.D., Smith, P., Grebel, D., Mahan, K., 1991. Haloperidol and nucleus accumbens dopamine depletion suppress lever pressing for food but increase free food consumption in a novel food choice procedure. Psychopharmacology 104(4), 515–521.

Salamone, J.D., Yohn, S.E., López-Cruz, L., San Miguel, N., Correa, M., 2016. Activational and effort-related aspects of motivation: neural mechanisms and implications for psychopathology. Brain 139(5), 1325–1347.

Schott, B.H., Niehaus, L., Wittmann, B.C., Schütze, H., Seidenbecher, C.I., Heinze, H.-J., Düzel, E., 2007. Ageing and early-stage Parkinson’s disease affect separable neural mechanisms of mesolimbic reward processing. Brain 130(9), 2412–2424.

Segar, T.M., Kasckow, J.W., Welge, J.A., Herman, J.P., 2009. Heterogeneity of neuroendocrine stress responses in aging rat strains. Physiology & Behavior 96(1), 6–11.

Seidler, R.D., Bernard, J.A., Burutolu, T.B., Fling, B.W., Gordon, M.T., Gwin, J.T., Kwak, Y., Lipps, D.B., 2010. Motor Control and Aging: Links to Age-Related Brain Structural, Functional, and Biochemical Effects. Neuroscience and biobehavioral reviews 34(5), 721–733.

Shephard, R.A., Broadhurst, P.L., 1982. Hyponeophagia and arousal in rats: Effects of diazepam, 5-methoxy-N,N-dimethyltryptamine, d-amphetamine and food deprivation. Psychopharmacology 78(4), 368–372.

Shoji, H., Miyakawa, T., 2019. Age-related behavioral changes from young to old age in male mice of a C57BL/6J strain maintained under a genetic stability program. Neuropsychopharmacology Reports 39(2), 100–118.

Sockeel, P., Dujardin, K., Devos, D., Denève, C., Destée, A., Defebvre, L., 2006. The Lille apathy rating scale (LARS), a new instrument for detecting and quantifying apathy: validation in Parkinson’s disease. J Neurol Neurosurg Psychiatry 77(5), 579–584.

Stuart, S.A., Hinchcliffe, J.K., Robinson, E.S.J., 2019. Evidence that neuropsychological deficits following early life adversity may underlie vulnerability to depression. Neuropsychopharmacology 44(9), 1623–1630.

Taylor, T.N., Greene, J.G., Miller, G.W., 2010. Behavioral phenotyping of mouse models of Parkinson’s disease. Behavioural brain research 211(1), 1–10.

Tombaugh, T.N., 2004. Trail Making Test A and B: Normative data stratified by age and education. Archives of Clinical Neuropsychology 19(2), 203–214.

Treadway, M.T., Buckholtz, J.W., Schwartzman, A.N., Lambert, W.E., Zald, D.H., 2009. Worth the ‘EEfRT’? The Effort Expenditure for Rewards Task as an Objective Measure of Motivation and Anhedonia. PLOS ONE 4(8), e6598.

Willner, P., 2016. The chronic mild stress (CMS) model of depression: History, evaluation and usage. Neurobiology of stress 6, 78–93.

Willner, P., Towell, A., Sampson, D., Sophokleous, S., Muscat, R., 1987. Reduction of sucrose preference by chronic unpredictable mild stress, and its restoration by a tricyclic antidepressant. Psychopharmacology 93(3), 358–364.

