## Supplementary methods for "Evidence for an apathy phenotype in aged mice"

The below includes a full description of the experimental method referenced in section 2.11 of the manuscript. This is a supplementary method for the website.

**Mass spectrometry analysis of corticosterone**

**Standard curve preparation**

Stock corticosterone and cortisol (Sigma Aldrich) were diluted in methanol to 100 ug/ml. 10 ul of each were diluted in 80 ul water to create a 10ug/ml working solution and then diluted in blood from an untreated c57bl/6j mouse to create a standard curve of the following concentrations: 0, 0.1, 0.3, 1, 3, 10, 30, 100 ng/ml. Corticosterone -D4 used to create the standard curve due to endogenous CS being present in the blood (response was the same as for authentic CS).

**Sample preparation**

When ready to assay Mitra devices were placed into 1.5ml screwtop eppendorf tubes containing 500ul of 0.5ngml D8- 17α hydroxyprogesterone (17α-OHP-D8 ) in 50:50 MeoH: Acn. Tubes were placed into a sonic bath for 15 minutes before being transferred to a vortex shaker set at 600rpm for 15 mins. Devices then removed from tubes. Samples were then blown to dryness under N_2_ in driblock heater at 40^o^C. Samples were reconstituted in 40ul for standards and spun at 13,000g for 5 minutes at room temperature. 35ul of spun supernatant was transferred to FIRV03 vials, capped and placed on an 8 °C CTC-HTS PAL autosampler (CTC analytics) for analysis.

**LC-MS/MS**

Following extraction HPLC-ES/MS-MS was performed using a Kinetex XB-C18 Column (6uM 100A 50 x 2.1mm) fitted with a 4 x 2.1mm C18 Guard cartridge from Phenomenex (Torrance, CA USA). The Column temperature was maintained at 40^o^C. The binary solvent system (2 Shimadzu LC-20ADXR pumps) utilised Milli Q H_2_O with 0.2mM ammonium Formate (eluent A) and Methanol (eluent B). The flow rate was 0.3ml/min with the following gradient 90%-A, 10% -B to 55% A, 45% B from 0 to 4 min, 45% -66%B from 4 to 7 mins, 66%-90%B from 7 to 8 mins held at 90% B from 8-12min returning to 10%B from 12 -13 mins and equilibrated at 10%B for a further 3 minutes giving a total run time of 16 minutes. MS detection was performed by an AB Sciex 5500 triple quad in positive electrospray (ESI+) The mass conditions were set as follows. Source/gas: Curtain Gas (CUR), 35psi; Collision Gas (CAD) 7 psi; Ion Spray Voltage (IS) 4500V; Temperature (TEM) 650^o^C; Ion Source Gas 1 (GS1) 65psi, Source Gas 2 (GS2) 75psi. Monitoring reaction Mode (MRM) was applied to the target compounds in quantitative analysis using Sciex Analyst 1.6.1 software to measure peak areas.

Results were determined by calculating a ratio of the internal standard (17a OHP-D8) to corticosterone peak area. A standard curve was then drawn in XLfit and sample ratios read against the curve to determine concentrations.

| ID | Q1 mass (Da) | Q3 mass (Da) | Dwell time (ms) | DP (v) | CE (v) | CXP (v) |
| --- | --- | --- | --- | --- | --- | --- |
| Corticosterone | 347.2 | 121.0 | 50 | 80 | 33 | 9 |
| Corticosterone-D4 | 351.2 | 121.0 | 50 | 86 | 33 | 10 |
| 17 αOHP-D8 | 339.2 | 100.0 | 50 | 71 | 31 | 12 |

**Table 1. Analytical parameters for corticosterone and 17a Hydroxyprogesterone-d8** Q1: precursor ion, Q3: product ion. DP: Declustering potential; CE: Collision energy; CXP: Collision exit potential.

The below includes a full description of the experimental method referenced in section 2.12-13 of the manuscript.

**c-Fos immunohistochemistry**

Brains were sliced in 40 µm coronal sections using a freezing microtome (Reichert, Austria). Sections were frozen in cryoprotectant (300 g sucrose, 300 ml ethylene glycol to 500 ml 0.1M phosphate buffer topped to 1L with dH_2_O) at – 20 ͦc until use.

Sections were washed 4 x 10 mins in TBS (Trizma buffer saline; 12g Trizma base, 9g NaCl, 1L dH_2_0 pH 7.4) and then incubated for 30 mins in TBS-T (Triton X, 0.1 %) with 3 % normal goat serum (Vector Laboratories) at room temperature. Sections were incubated with primary antibody rabbit anti c-Fos (ABE457, Merck Millipore) 1:4000 in TBS-T with 3 % normal goat serum overnight in the fridge at 4 ͦc. The following day sections were washed in TBS-T with 3 % normal goat serum. Sections were then incubated with secondary antibody goat anti-rabbit (1:500, Alexa Fluor 488, Invitrogen) in TBS-T with 3 % normal goat serum for 3 hours. The following steps were done in the dark. Sections were washed 3 x 10 mins in TBS. Sections were then incubated with DAPI (14.3 µM) for 3 minutes and then washed in PBS 3 x 5 minutes. Sections were then mounted onto subbed slides (VWR) and a few drops of Vectashield mounting media was added (Vector Laboratories). They were then cover-slipped and stored in the fridge in darkness until imaging.

Images were taken on a Leica widefield microscope with DFC365 FX camera with LasX software. For images taken with the 10x objective, the cfos channel had a 500 ms exposure and 2.7 gain while the DAPI channel had 100 ms exposure. For images taken with the 5x objective the cfos channel had 999 ms exposure and 2 gain while DAPI had a 500 ms max exposure. 2 brain regions were captured; paraventricular nucleus of the hypothalamus (PVN) at using the 10x objective and amygdala (central and basolateral regions) using the 5x objective. A picture of the whole PVN was taken across 3 different sections, bregma levels ranging from -0.58 to -0.94. A picture of the amygdala (left or right) was taken across 3 different sections, bregma levels ranging from -1.22 to -1.40. Some brains/sections were lost due to tissue damage. N = 10 per group amygdala was obtained. N = 10 PVN for aged and young group were obtained, N = 9 for middle aged group.

Cfos was counted manually using ImageJ cell counter. The brain region of interest was manually drawn around and cfos was counted within that area. Contrast was adjusted uniformly across all images. Cfos count was normalised by dividing cfos count by area. C-fos dots were only counted if they co-localised with DAPI. Experimenter was blind to age group.
